# Rieske Iron-Sulfur Cluster Proteins from an Anaerobic Ammonium Oxidizer Suggest Unusual Energetics in their Parent Rieske/cytochrome *b* complexes

**DOI:** 10.1101/2025.06.25.661457

**Authors:** David Hauser, Mandy Sode, Elena A. Andreeva, Kristian Parey, Thomas R.M. Barends

## Abstract

Anaerobic ammonium-oxidizing (anammox) bacteria employ a unique, hydrazine-based pathway to obtain energy from nitrite and ammonium. These organisms express distinct Rieske/ cytochrome *b* complexes whose precise role in anammox metabolism remains unclear, but which has been proposed to include the generation of NAD(P)H. This would require energetics and structural features unusual for such complexes. Here we present crystal structures and electrochemical investigations of the Rieske subunits of two of these complexes from the anammox organism *Kuenenia stuttgartiensis*, Kuste4569 and Kustd1480. Both proteins display high redox potentials (>+300 mV), which can be in part explained by their crystal structures and which fit perfectly into the energetic scheme of the proposed NAD(P)H generation mechanism. Moreover, AlphaFold3 models of the parent complexes trace out a path for the electrons required for NAD(P) production, which includes a proposed, novel *b*-type heme in the membrane-bound part of the complex.

## Introduction

Anaerobic ammonium oxidation (“anammox”)^1,2^ is a bacterial pathway in which ammonium is condensed with nitrite to yield nitrogen gas and water, providing energy for the cell. The anammox process, which was only discovered in the 1990s, is of global importance: it is responsible for up to 10% of the global nitrogen cycle and, in some ecosystems, contributes up to 50% to the total yearly N_2_ production. Anammox metabolism consists of a set of redox reactions taking place in a dedicated compartment known as the anammoxosome (Figure 1a). In a first step, nitrite (NO ^-^) is converted to nitric oxide (NO), which is then condensed with ammonia (NH_3_) to generate the extremely unusual, highly reactive compound hydrazine (N_2_H_4_). The hydrazine is subsequently oxidized to dinitrogen gas (N_2_), releasing four electrons of extremely low redox potential. According to our current understanding of the anammox process ^1,2^, these are believed to be passed on to an as yet unidentified membrane protein, which uses them to pump protons into the anammoxosome. The proton gradient thus produced is used by one or more ATPases to synthesize ATP. Moreover, this unknown membrane protein has been hypothesized to also reduce lipoquinone, which was proposed to then be used by one or more Rieske/cytochrome *b* complexes to pump even more protons over the anammoxosome’s membrane.

**Figure 1.**
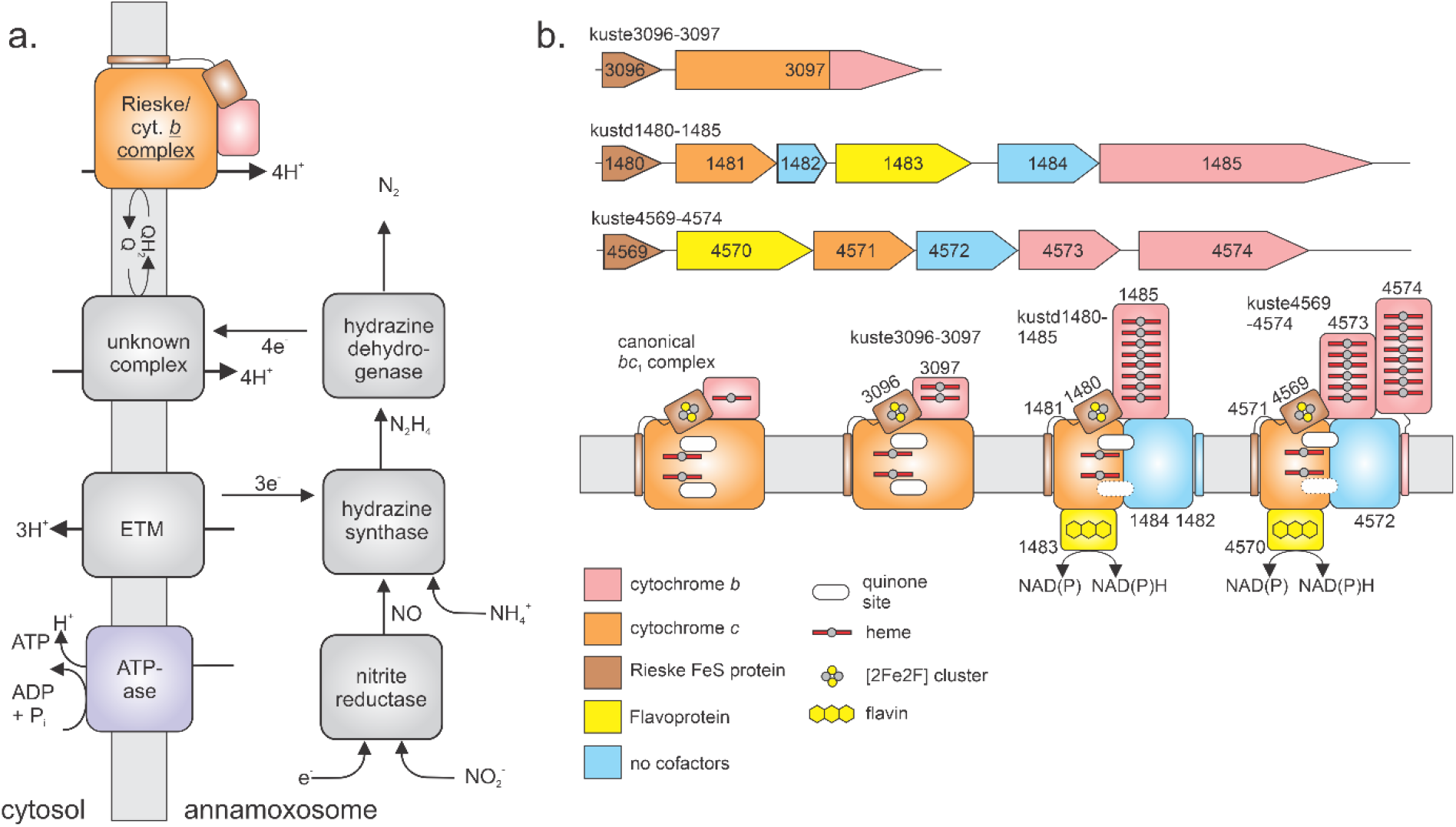
**a**. The anammox process involves a series of redox reactions driving an electron transfer chain used to pump protons into the anammoxosome. The resulting proton gradient is then used to fuel ATP synthesis. **b**. Gene- and proposed domain organization of Rieske/cytochrome *b* complexes in *K. stuttgartiensis* as was presented by Kartal *et al*.^*1*^.

Rieske/cytochrome *b* complexes^3 4 5^ are found in most organisms that use chemiosmosis to produce a proton gradient to drive ATP synthesis. In general, there are two major classes of these complexes. The first is made up by the *bc*_*1*_ complexes found in many prokaryotes and in mitochondria, where they are also referred to as “Complex III” ^3^. The second class are the *b*_*6*_*f* complexes ^4^ found in organisms performing oxygenic photosynthesis, where they are mediating electron transfer between PSII and PSI ^6^. Both types of complexes contain a membrane-bound cytochrome and a “Rieske iron-sulfur protein”, which consists of a small domain containing a [2Fe2S] cluster, typically connected to a single transmembrane helix. Importantly, Rieske/cytochrome *b* complexes make use of *electron bifurcation* to harvest the energy released by the two-electron oxidation of a lipoquinone as efficiently as possible, while coupling it to the reduction of a soluble one-electron transfer protein with a considerably higher potential. The process involves the oxidation of a quinone molecule at a site close to the positive side of the membrane (called the “Q_P_” site), which results in the release of two protons. The two electrons resulting from this oxidation then follow different paths. One electron follows an energetically favorable route, being first transferred to the high-potential iron-sulfur cluster in the Rieske protein. The Rieske protein then reorients and transfers the electron to the *c*-type cytochrome domain of the complex and onward to a soluble redox partner. This “downhill”, energetically favorable transfer enables the other electron to take an “uphill”, energetically unfavorable route, in which it is transferred to a low-potential heme of the cytochrome *b* component. From there, the electron can be transferred to another quinone binding site close to the negative side of the membrane (the Q_N_ site) where it performs the one-electron reduction of a quinone to a semiquinone radical. This is followed by another cycle of quinone oxidation and electron bifurcation, with the resulting electrons again following different paths, one going favorably *via* the Rieske protein to the *c-*type cytochrome and from there to another molecule of the soluble redox partner, and the other electron again undergoing the unfavorable transfer to the low potential heme in the *b* component. From there, it is used to further reduce the semiquinone to a fully reduced lipoquinone which can re-enter the quinone pool. In the entire process, two quinone molecules are thus oxidized in the Q_P_ site, releasing four protons at the positive side of the membrane, whereas one quinone molecule is reduced at the Q_N_ site, resulting in two protons being taken up at the negative side of the membrane. This elegantly couples the two electron transfer cycles to proton translocation.

Most organisms require only a single Rieske/cytochrome *b* complex for their energy metabolism. Strikingly however, the anammox model organism *Kuenenia stuttgartiensis* encodes a total of three such complexes on its genome (Figure 1b). Other species have been found that also contain more than one copy of Rieske/cytochrome *b* complexes, although generally little is known about their energy metabolisms^7^. In proteomics studies, all three of the complexes of *K. stuttgartiensis* have been found to be expressed^8^. The simplest of the *K. stuttgartiensis* Rieske/cytochrome *b* complexes in terms of gene structure, Kuste3096-3097 (named after their gene loci), appears to be the most similar to a canonical *bc*_*1*_ complex, with two notable differences: first, the equivalent of cytochrome *c*_1_ contains two hemes and second, this domain is fused with the cytochrome *b* into a single polypeptide chain (as is also seen often in gram-positive bacteria^5,9^). The other two complexes, Kustd1480-1485 and Kuste4569-4574, the latter being the most highly expressed of the three, have more peculiarities, and seem to be more closely related to *b*_*6*_*f* rather than to *bc*_*1*_ complexes. Indeed, both feature cytochrome *b*_*6*_-like as well as “subunit IV(SUIV)”-like transmembrane proteins^1^. However, instead of cytochrome *f*, these complexes contain a hexa- or octaheme cytochrome *c* and in the case of Kuste4569-4574, an additional HAO-like octaheme cytochrome *c* (OCC) is present. Most strikingly however, both complexes contain a putative NAD(P) oxidoreductase, likely placed on the riboplasmic side of the membrane^1^.

Given the presence of these unexpected NAD(P) oxidoreductases in two of the three Rieske/cytochrome *b* complexes in *Kuenenia*, a novel function and mechanism for both these complexes has been proposed^1^, in which the electron bifurcation process could be used for the production of NAD(P)H. In this proposal, one of the two electrons from quinol oxidation would not be used for the reduction of another quinone, but for the energetically unfavorable reduction of NAD(P)+ to NAD(P)H 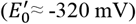, whereas the second electron is used for another, energetically favorable reaction. In the case of the Kuste4569-4574 complex, the presence of the HAO-like OCC suggests that this reaction could be the reduction of nitrite to NO 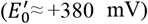 ^1^. Overall, this coupling of NAD(P)+ reduction to nitrite reduction should be feasible thermodynamically, since the most abundant quinone species in *K. stuttgartiensis* has been shown to be menaquinone-7 (MK-7) ^8^ which has an 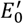 of about -70 mV ^10^. In the Kustd1480-1485 complex the situation is similar; however, no proposal has been made as to what energetically favorable redox reaction could be used to drive NAD(P) reduction.

In this study, we focus on the Rieske iron-sulfur cluster proteins of the two NAD(P) reductase-containing Rieske/cytochrome *b* complexes from *K. stuttgartiensis*. We study their crystal structures and redox potentials, and discuss these in the context of the thermodynamic constraints of the proposed reactions, supported by AlphaFold3 models of the respective complexes as a whole.

## Materials and Methods

### Protein production

Vectors based on pET24d(+) were designed for expression of the soluble domains (*i*.*e*., excluding the transmembrane helices and their linkers to the soluble parts) of Kustd4569 and Kustd1480 guided by AlphaFold3 ^11^ and ordered from GenScript B.V. (Rijswijk, the Netherlands). Proteins were expressed in *E. coli* BL21 suf++. Cells were cultured in 2 L LB medium supplemented with 50 μg/mL kanamycin sulfate in 5L baffled flasks. These were inoculated with 20 mL of preculture and grown at 37°C, 120 rpm until the OD_600_ reached 0.4-0.6. Temperature and shaker speed were then reduced to 18°C and 90 rpm, respectively, and IPTG, L-cysteine and ammonium iron-(III) citrate were added to final concentrations of 1 mM each. The cultures were harvested after 20 h by centrifugation with a Sorvall Lynx 6000 centrifuge equipped with an F9-6×1000 LEX rotor (ThermoFisher Scientific, Bremen, Germany) at 4°C, 6000 rpm, 15 min. Cell pellets were resuspended in wash buffer (300 mM sodium chloride, 50 mM Tris/Cl pH 8.0, 10 mM imidazole), frozen in liquid nitrogen and stored at -20°C. For purification, thawed cell suspensions were supplemented with *Complete™ EDTA-free* protease inhibitor cocktail and incubated for up to 30 minutes on ice. Cells were then lysed by sonication using a W-450 D Branson Digital Sonifier at 30% amplitude and 1 s cycles for 8 min, and the lysate centrifuged using a Sorvall Lynx 6000, in an F20-12×50 LEX rotor (ThermoFisher Scientific, Bremen, Germany) at 20,000 rpm, 4°C, for 1 h. The cleared lysate was filtered through 0.45 μm syringe filter and loaded onto a column with 4 mL Ni-NTA beads (Qiagen, Hilden, Germany) pre-equilibrated with wash buffer. The column was washed with 3×15 mL wash buffer, and bound proteins were eluted with ∼5 mL of wash buffer with 250 mM imidazole. The eluted protein was dialyzed overnight against wash buffer without imidazole overnight using 3.5 kDa RC Tubing (Spectrumlabs, Piraeus, Greece). The proteins were then concentrated to a volume of 1 mL and loaded onto a Superdex S75 16/600 GL (GE Healthcare, Uppsala, Sweden) column, equilibrated in gel filtration buffer (150 mM sodium chloride, 50 mM Tris pH 8). Flow rates of 0.5-1.5 mL/min were used and 1 mL fractions collected. Colored fractions were investigated by SDS-PAGE and sufficiently pure fractions combined. After concentrating to 0.5-1 mL, 50 μL aliquots were prepared frozen in liquid nitrogen and stored at -80°C. All proteins were stored in gel filtration buffer. Protein identities were confirmed by peptide mass fingerprinting.

### UV/Vis spectroscopy

Proteins were diluted in gel filtration buffer and UV/Vis spectra measured between 250 and 800 nm (0.5 nm bandwidth, scan speed 400 nm/min) in quartz cuvettes with a 1.0 cm path length (Hellma GmbH, Müllheim, Germany) using a JASCO V-650 spectrophotometer (Jasco GmbH, Gross-Umstadt, Germany) at room temperature. To collect spectra of oxidized proteins, samples were diluted in gel filtration buffer containing 1 mM potassium ferricyanide, and the spectrometer blanked against the same solution without protein prior to data collection. Spectra of protein thus diluted in ferricyanide-containing buffer were collected after 1- and 90 minutes intervals and compared, with no differences observed.

### Spectroelectrochemistry

Spectroelectrochemical experiments were carried out in a custom-built optically transparent thin-layer electrochemical (OTTLE) cell ^12^ with a gold mesh electrode freshly modified with a 4,4’-dithiopyridine solution (20 mM in 160 mM Tris-Cl, 20% ethanol (v/v)) for at least 1 hour at room temperature. The cell was connected to a Keithley model 2450 source measure unit (Tektronix Inc., Beaverton, USA) running as a potentiostat to perform chronoamperometry and initiate the measurement of spectra. Every sample was pre-poised at the first potential for 10 min. UV/Vis spectra were recorded in 20-50 mV potential steps using a Jasco V-760 Spectrophotometer (Jasco GmbH, Gross-Umstadt, Germany). The sample (in 10mM MOPS/KOH pH7, 100mM KCl) was allowed to equilibrate for 2 min at every potential before collecting a spectrum. The current range of the source measure unit was limited to 1 mA.

All spectra were baseline-corrected using the absorbance averaged between 700 nm and 650 nm to increase the signal to noise ratio. The baseline-corrected absorbance at wavelengths of maximal change (465-485 nm for both Kuste4569 and Kustd1480) was then extracted, averaged over the wavelength range and normalized. However, due to the nature of the cell, the path length is not completely fixed and can decrease during the measurement due to capillary forces in the parafilm sealing the cell and/or evaporation. For this reason, a linear correction was applied based on the absorbance difference of the first and the last spectrum that were taken at the same potential, which, importantly, has little influence on the fit results but improves plot visuals. The corrected, normalized absorbance was then plotted against the potential and fitted to a single-transition Nernstian function using a custom-written Python script calling NumPy ^13^, Pandas ^14^, Scipy ^15^ and Matplotlib ^16^:

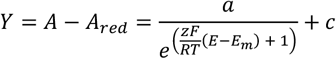

where *A* is the averaged and normalized absorbance, *A*_*red*_ the absorbance of the assumed fully reduced reference spectrum taken at the lowest potential, *a* the amplitude of the transition, *z* the number of electrons (set to 1 in this case), *E* the potential set by the SMU, *E*_*m*_ the midpoint potential and *c* an offset. The Faraday constant *F* was taken to be 96,485.34 J mol^-1^ V^-1^, the ideal gas constant *R* 8.3145 J mol^-1^ K^-1^ and the temperature *T* was 293 K. The offset of the Ag/AgCl reference electrode (a thin patch of Ag/AgCl ink) at 100 mM KCl was taken to be +280 mV against the standard hydrogen electrode (SHE) based on comparison with a commercial Ag/AgCl electrode (Pine Research, Durham, USA) and confirmed by spectropotentiometric measurements on bovine cytochrome *c*.

The error of the fitted midpoint potential was determined using parametric bootstrapping assuming a Gaussian distribution. Using a custom-written Python script, 1000 data sets were generated based on the experimentally determined means and Gaussian noise added to the data using the experimental standard deviations. The standard deviations were scaled by a factor of 3 before generating noise. Nernstian fits were applied to every bootstrapped dataset and the standard deviation of the resulting distribution of midpoint potentials was taken as the estimate for the variation in the midpoint potential of the 3 experimental replicates.

### Protein Crystallization and Structure Determination

To find initial crystallization conditions, the commercial screens of the JCSG Core Suite and Ammonium Sulfate Suite from NeXtal Biotechnologies (Holland, USA) were used. Using a Mosquito robot (SPT Labtech Ltd., Melbourn, UK), 100 nL of concentrated protein solution were mixed with 100 nL of reservoir solution in 96-well sitting-drop plates (Greiner XTL low-profile, Greiner Bio One, Frickenhausen, Germany). Conditions were optimized to obtain larger crystals using sitting drops of 1 μL protein mixed with 1 μL of reservoir solution with glass slides placed on 24-well plates (Crystalgen SuperClear CPL-132, Jena Bioscience GmbH, Jena, Germany) filled with 700 μL of reservoir solution per well.

Raw x-ray diffraction data were processed using XDS ^17^. Both structures were solved by molecular replacement using PHASER ^18^ with AlphaFold 3 models ^11^ as the search models. The initial models went through iterative rounds of refinement in PHENIX ^19^ and manual correction in COOT ^20^. A lattice translocation defect in the Kuste4569 dataset was corrected for using the method of Wang *et al*.^21^ using the programs ‘SFTOOLS’ and ‘FFT’ of the CCP4 suite ^22^ to edit the reflection data in the mtz file and to calculate difference Patterson maps ^22^. Further details are given in the Supplemental Information. Ensemble refinement for Kustd1480 was performed with PHENIX as described in ^23,24^. A three-dimensional grid search was set up, scanning different values for *p*_TLS_, *T*_bath_ and *t*_x_. The individual refinements were carried out in parallel on 36 cores of a cluster of Intel^**®**^ Xeon^**®**^ CPU X7560 processors running at 2.3 GHz. The structure with the lowest difference between R_free_ and R_work_ was selected as the final ensemble model. Structural analysis was performed using COOT/PyMOL ^25^ and custom-written python scripts calling NumPy and Matplotlib. Figures were prepared using PyMOL.

### Modeling of complexes

Models of the full Kustd1480-1485 and Kuste4569-4574 complexes were generated using AlphaFold3^11^, running on the AlphaFold server (www.alphafoldserver.com). All expected hemes *b* and *c*, as well as one novel heme *b* were included by the server, as were NADP and FAD. The only difference between a *b*- and *c*-type heme is that the latter is covalently attached to least one cysteine *via* one or both vinyl group(s). We observed that while AlphaFold3 places heme groups in their expected positions given the locations of heme binding domains, and predicts the covalent attachments in the case of *c*-type heme, it often labels these as *b*-type hemes and *vice versa* regardless of binding mode. Assignment of heme types were therefore done manually. The [2Fe2S] clusters from the Rieske proteins were added manually based on alignments of the crystal structures of these proteins to the AF3 models. Further iron-sulfur clusters were placed in the putative NAD(P) reductases Kustd1483 and Kustd4570 as follows: the predicted fold of Kustd4570 was submitted to a search for structural homologs using the DALI server ^26^. This identified domains of the Bfu family member NfnABC from *Caldicellulosiruptor saccharolyticus* (pdb entry 9bp5 ^27^) and of the tungsten-containing formylmethanofuran dehydrogenase from *Methanothermobacter wolfeii* (pdb entry 5t5i ^28^) as having structural homology (RMSD 2.7 Å for 483 amino acids and 4.2 Å for 335 amino acids, respectively). These experimental structures contain multiple [4Fe4S] iron-sulfur clusters which aligned excellently with groupings of cysteine residues in Kustd1483 and Kustd4570, and these clusters from the aligned homologs were therefore used to complete the AF3 model. Due to the strong structural homology of Kuste4570 to Kustd1483, these clusters could also be used to complete the model of the latter protein. In both proteins, however, the iron-sulfur cluster closest to the FAD lacks a fourth coordinating cysteine, and a buried lysine closely approaches the cluster. This cluster was therefore modeled as an [3Fe4S] cluster. Initial, partial models also included one chlorophyll molecule per monomer, as seen in *b*_6_*f* complexes, but these cofactors were omitted in the final models to save computational expenses and because no chlorophyll synthetic machinery seems to be present in *K. stuttgartiensis*.

## Results

### Crystal structure determination of Kustd1480 and Kuste4569

Expression constructs for the soluble domains of the Rieske iron-sulfur cluster proteins Kustd1480 and Kuste4569, *i*.*e*., without their respective transmembrane helices and linkers, were designed and the proteins expressed heterologously. Both proteins were purified using metal ion affinity- and size exclusion chromatography. Kustd1480 was concentrated to 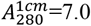, and crystallized in 1+1 μL hanging drops equilibrated using 0.1 M CHES pH 9.5, 17% PEG 8000 as the reservoir solution. Reddish brown, lozenge-shaped plates grew within one day and were cryocooled in liquid nitrogen after cryoprotection in mother liquor supplemented with 20% (v/v) glycerol. Data were collected to 1.8 Å resolution but showed severe anisotropy. These data were phased successfully using an AlphaFold3 model as the search model (TFZ=12.8 without cofactors in the model), but upon refinement, the R-factors did not decrease as expected. Inspection of the data and the molecular replacement solution revealed strong noncrystallographic translation symmetry, which often hampers refinement as it causes correlations in the intensities that are not accounted for in the model^29,30^. Moreover, it can artificially increase the R-factors, particularly at high resolution, as a considerable fraction of the reflections will be very weak. We therefore cut the resolution to 2.5 Å, but while this improved the R-factors considerably, they never reached values below 30%. Inspection of the density revealed severe smearing out of the data in various distinct directions in several parts of the structure, such as around the iron sulfur clusters. After testing various possible explanations, such as incorrect space group assignment with or without various forms of twinning, it was found that the most parsimonious explanation was structural disorder in the protein. Indeed, the regions affected are identical in both molecules in the asymmetric unit, with the relative direction of the apparent disorder being the same in both molecules for each region, which is consistent with it being a property of the protein rather than a crystallographic artefact. To account for this disorder, we performed ensemble refinement, which resulted in an ensemble model of good R-factors and acceptable geometry (see Table 1 for data and refinement statistics).

**Table 1.** Data collection and refinement statistics.

|  | **Kustd1480**<br>**pdb entry 9rk3** | **Kustd4579**<br>**pdb entry 9rk4** |
| --- | --- | --- |
| **Data collection** |  |  |
| Space group | *P*2<sub>1</sub>2<sub>1</sub>2<sub>1</sub> | *P*2<sub>1</sub>2<sub>1</sub>2<sub>1</sub> |
| a, b, c (Å) | 46.2, 76.2, 79.6 | 104.0, 120.7, 209.2 |
| $\alpha$, $\beta$, $\gamma$ (°) | 90, 90, 90 | 90, 90, 90 |
| Wavelength (Å) | 0.88560 | 0.88560 |
| Resolution range (Å) | 40-1.80 (1.85-1.80) | 104.6 – 2.8 (2.9 – 2.8) |
| Total no. of reflections | 86,269 (6,681) | 879,981 (83,186) |
| No. of unique reflections | 25,291 (1,871) | 65,436 (6,430) |
| Completeness (%) | 94.4 (95.3) | 99.9 (100.0) |
| Redundancy | 3.4 (3.6) | 13.4 (12.9) |
| $\langle I/\sigma(I) \rangle$ | 8.2 (1.9) | 13.0 (1.9) |
| $R_{\text{merge}}$ | 0.065 (0.729) | 0.136 (1.622) |
| $CC_{1/2}$ | 0.998 (0.909) | 0.999 (0.803) |
| Wilson B-factor (Å<sup>2</sup>) | 33.8 | 64.9 |
| **Refinement** |  |  |
| Resolution range (Å) | 40-2.5 | 73.7 – 2.8 |
| Completeness (%) | 93.6 | 99.5 |
| No. of reflections | 9,570 | 65,248 |
| Final $R_{\text{cryst}}$ | 0.2205 | 0.2645 |
| Final $R_{\text{free}}$ | 0.2761 | 0.2972 |
| No. of non-H atoms |  |  |
| Protein | 14,848: 8 models, 2 monomers/AU | 10,260: 12 monomers/AU |
| Ligands | 64 [2Fe2S], 8 models | 48 [2Fe2S], 60 SO<sub>4</sub><sup>2-</sup> |
| Water | 185, 8 models | n.a. |
| R.m.s. deviations |  |  |
| Bonds (Å) | 0.019 | 0.002 |
| Angles (°) | 1.71 | 0.515 |
| Average *B* factors (Å<sup>2</sup>) |  |  |
| Protein | 47.4 | 72.5 |
| Ligands | 47.7 | 72.4 |
| Water | 45.7 | n.a. |
| Ramachandran plot |  |  |
| Most favoured (%) | 86.3* | 96.5 |
| Allowed (%) | 10.5* | 3.4 |
| Outliers | 3.3* | 0.2 |
| Rotamer outliers (%) | 12.8* | 1.8 |

\* A large number of unfavorable ( $\phi$ , $\psi$ ) combinations and rotamers is expected, as ensemble refinement samples high-energy conformers<sup>24</sup>

Kuste4569 was concentrated to an 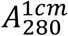 of 12 and crystallized in 1+1 μl hanging drops using 4% (v/v) 2,2,2-trifluoroethanol, 2 M ammonium sulfate as the reservoir solution. Brown, lozenge shaped crystals grew within one week, which were cryoprotected in mother liquor with 20% (v/v) glycerol before cryocooling in liquid nitrogen and data collection. Phasing of the 2.8 Å resolution data was done by molecular replacement using an AlphaFold3 model as the search structure (TFZ=22), however, upon refinement the R-factors failed to converge for this structure as well. As described in detail in the Supplemental Information, this was found to be due to a lattice translocation defect, which was corrected at the intensity level using the method of Wang *et al*. ^21^. This allowed the structure to be refined to good geometry and R-factors, as reported in Table 1.

The soluble domains of both Kustd1480 and Kuste4569 display the expected fold of a Rieske iron-sulfur protein, consisting of a cofactor binding domain with an incomplete β-barrel topology that is an insertion on a larger, “basal” domain as shown in Figure 2. Indeed, the structures of Kuste4569 and Kustd1480 are very similar (Figure 2, Supplemental Figure S4), with an RMSD of 1.3 Å for 106 C_α_ atoms, consistent with the generally high structural conservation among Rieske proteins. Both proteins coordinate their [2Fe2S] cluster with the canonical motif of two conserved histidines and two cysteines (Figure 3, Figure 4). The structures can also be aligned well with Rieske proteins from other species. A search for homologous structures of Kuste4569 using the DALI server ^26,31^ showed it to be most similar to the Rieske protein from the *Thermosynechococcus elongatus* BP-1 *b*_6_*f* complex (pdb entry 3azc) ^32^. Other hits with high structural similarity were Rieske proteins from other *b*_6_*f* complexes, as well as Rieske-type proteins from arsenite oxidases. Small differences are found in the cluster binding domains and the other two β-sheets, while the loops frequently show insertions of various sizes, as has been noted before ^33,34^. However, the C-termini of Kuste4569 and particularly Kustd1480 appear to be longer than in many other Rieske proteins; few examples, such as the *Spinacia oleracea* protein(Figure 4) feature a tail of comparable length. The long C-terminus of the *S. oleracea* protein appears to be typical for Rieske proteins in *b*_6_*f* complexes ^32^. However, unlike those of the *b*_6_*f* complex Rieske proteins, the ends of the C-terminal tails of Kuste4569 and Kustd1480 are hydrogen-bonded to β-strand 4 for a short stretch in a β-sheet pattern at the very end. The C-terminal part of Kustd1480, which is the longest of the two, forms a loop on the surface of the protein (Figure 2, Supplemental Figure S4). The flexibility observed in the Kustd1480 structure mostly concerns this C-terminal loop, as well as the loop between β3 and β4 (which link the basal- and cofactor binding domains) and part of the “proline loop” (which covers one side of the cofactor, and whose C-terminal part forms the second link between the basal- and cofactor binding domains) (Supplemental Figure S5), but also the relative position of the cofactor-binding domain, which appears to be moving as a rigid body with respect to the basal domain. The amplitude of this domain’s flexibility is largest at its outermost tip, which appears to move from side to side by as much as 2 Å (Figure 2).

**Figure 2.**
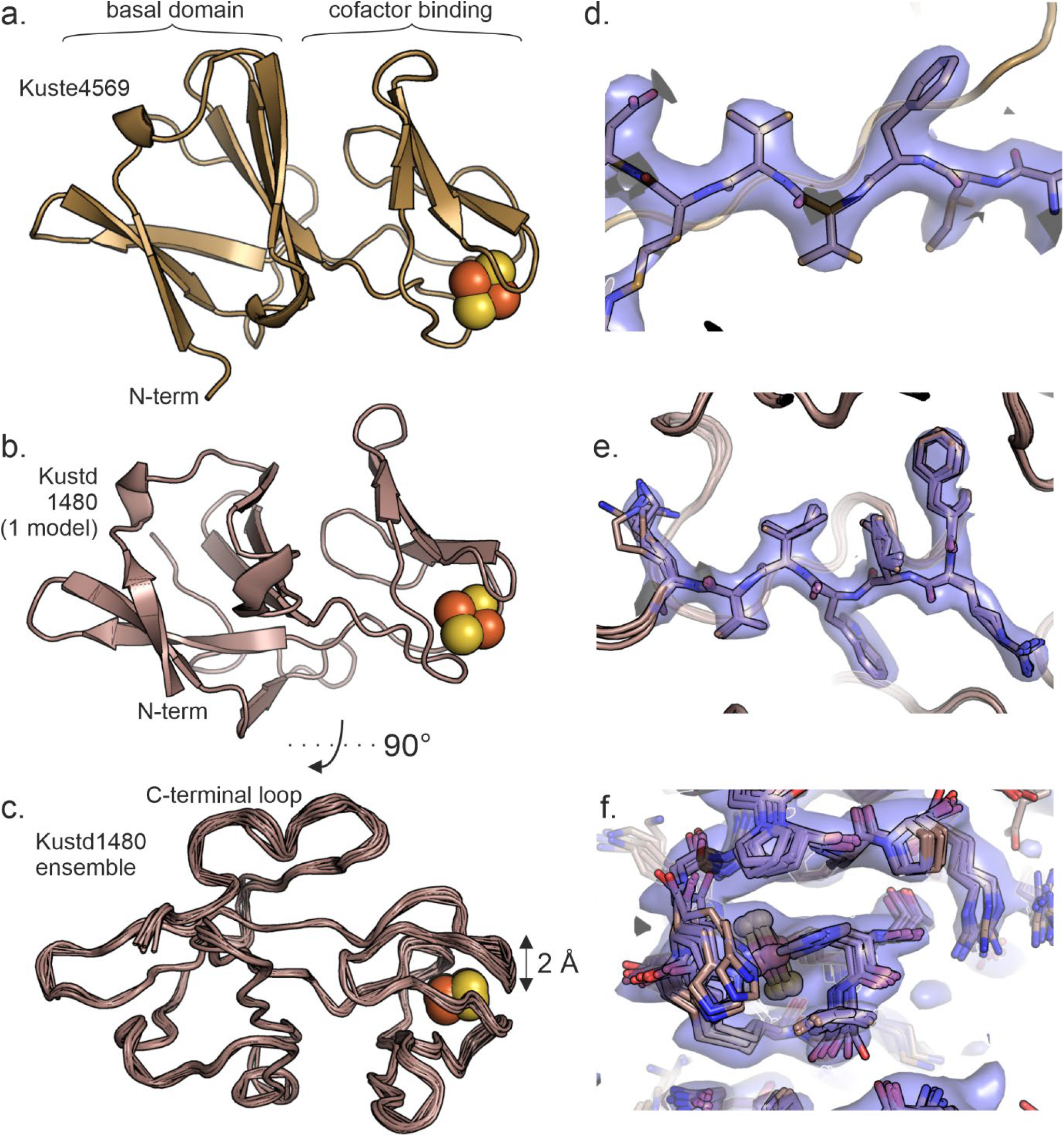
Crystal structures of the soluble part of **a**. Kuste4569 and **b**. Kustd1480. The [2Fe2S] cofactors are shown as yellow and orange spheres. **c**. Ensemble structure of Kustd1480. The largest degree of flexibility is seen around the cofactor and in the C-terminal loop. **d**. Close up of a portion of the 2*m*Fo-*D*Fc electron density map for Kuste4569. **e**. Close-up view of part of the 2*m*Fo-*D*Fc electron density map for Kustd1480 in a well-ordered part of the structure, and **f**. in the cofactor binding domain. The latter shows considerable disorder in one direction. All maps were contoured at 1 σ and shown together with the final, refined structure (the ensemble structure in the case of Kustd1480).

**Figure 3.**
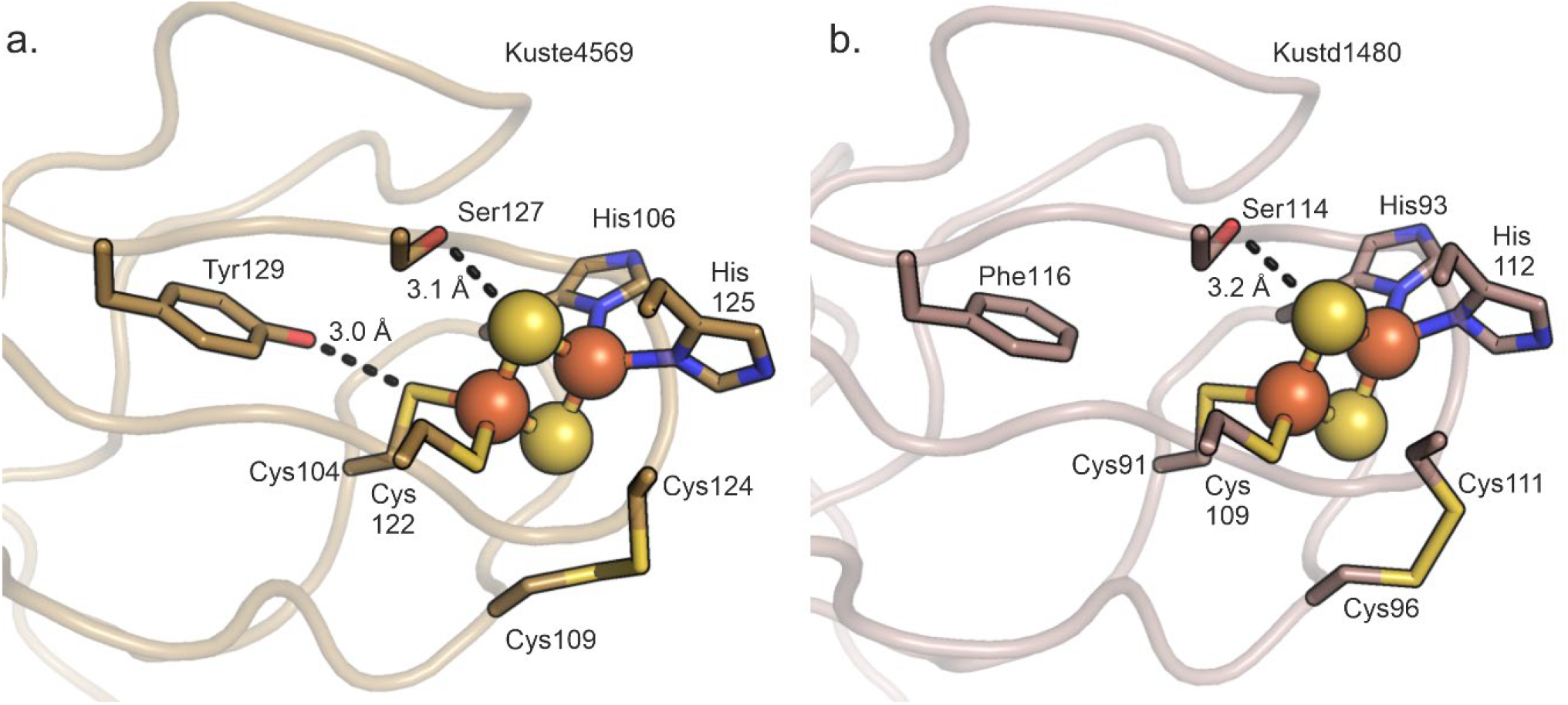
Coordination and surroundings of the [2Fe2S]-cofactors in. **a**. Kuste4569 and **b**. Kustd1480. The [2Fe2S] clusters are shown as balls and sticks. The iron-coordinating histidines and -cysteines are shown as sticks, along with the disulfide bridge and the Tyr129/Ser127-pair (Phe116/Ser114 for Kustd1480) implicated in redox potential tuning^10,36,37^. Interaction distances are indicated in Å.

**Figure 4.**
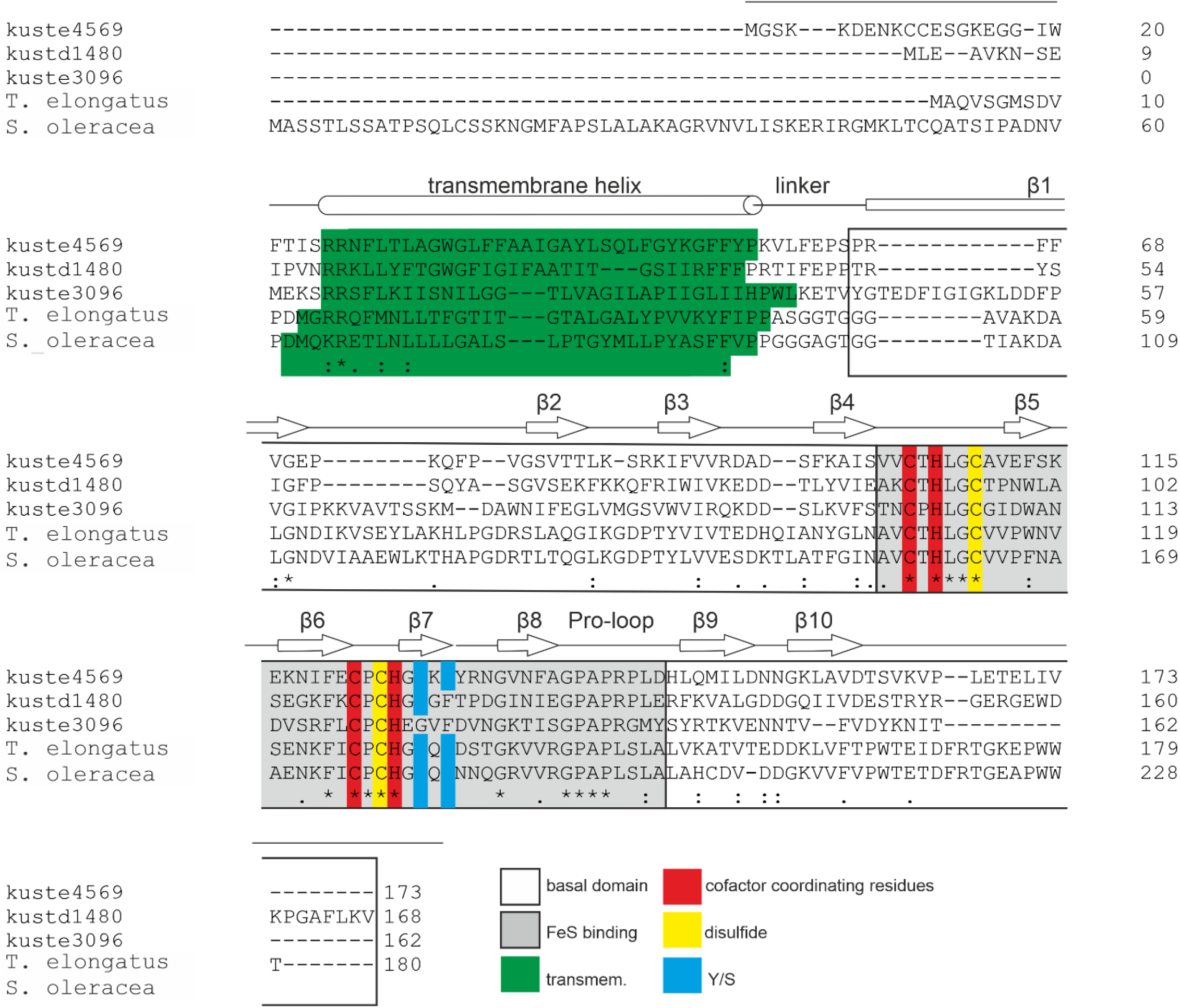
Multiple sequence alignment of Rieske proteins from *Kuenenia stuttgartiensis* (kuste4569, kustd1480, kustd3096) and the closest structural homologs of kuste4569, the Rieske proteins from *Thermosynechococcus elongatus* (WP_011056803.1) and *Spinacia oleracea* (NP_001413377.1). Secondary structure elements are indicated for the soluble domain of Kuste4569 according to its crystal structure; the positions of the transmembrane helices were predicted using AlphaFold3 ^11^. The basal- and cofactor-binding domains are indicated as are the positions of the binding domain’s disulfide bridge, the cofactor coordinating residues, and the Tyr/Ser pair (“Y/S”) implicated in tuning the redox potential^10,36,37^.

The high redox potential of Rieske proteins in comparison to other [2Fe2S]-type iron-sulfur cluster proteins is not only due to their clusters being coordinated by two histidines and two cysteines instead of four cysteines, but also due to residues in the second coordination sphere, in particular the hydrogen bonding network around the cluster ^35^. Here, two positions in the sequence at the end of strand β7 appear to be of particular importance (Figure 4); presence of a tyrosine and a serine in these positions increases the redox potential by hydrogen bonding to a cofactor-coordinating cysteine- and a cluster sulfur, respectively^10,36,37^. In Kuste4569, both these residues (Tyr129/Ser127) are present and engage in the expected interactions.

In contrast, Kustd1480 has a phenylalanine (Phe116) in the position of the tyrosine, although the serine (Ser114) is present and interacts with the cluster (Figure 3).

### Spectropotentiometry

In their as-isolated states, both proteins show similar UV/Vis spectra (Figure 5a,b), apart from a large difference in absorption around 280 nm, which is much higher in the case of Kustd1480 than for Kuste4569. This is easily explained by the absence of tryptophan residues in the latter protein. Moreover, in its oxidized form Kustd1480 has an absorption peak at 430 nm, whereas Kuste4569 has no peak there in the as-isolated state, which may contribute to the somewhat more reddish hue of Kustd1480 in solution. Upon oxidation, both proteins show increases in absorption over a broad range of absorptions, particularly between 320 and 380 nm and between 420 and 580 nm. These changes were used for spectropotentiometric redox potential determination.

**Figure 5.**
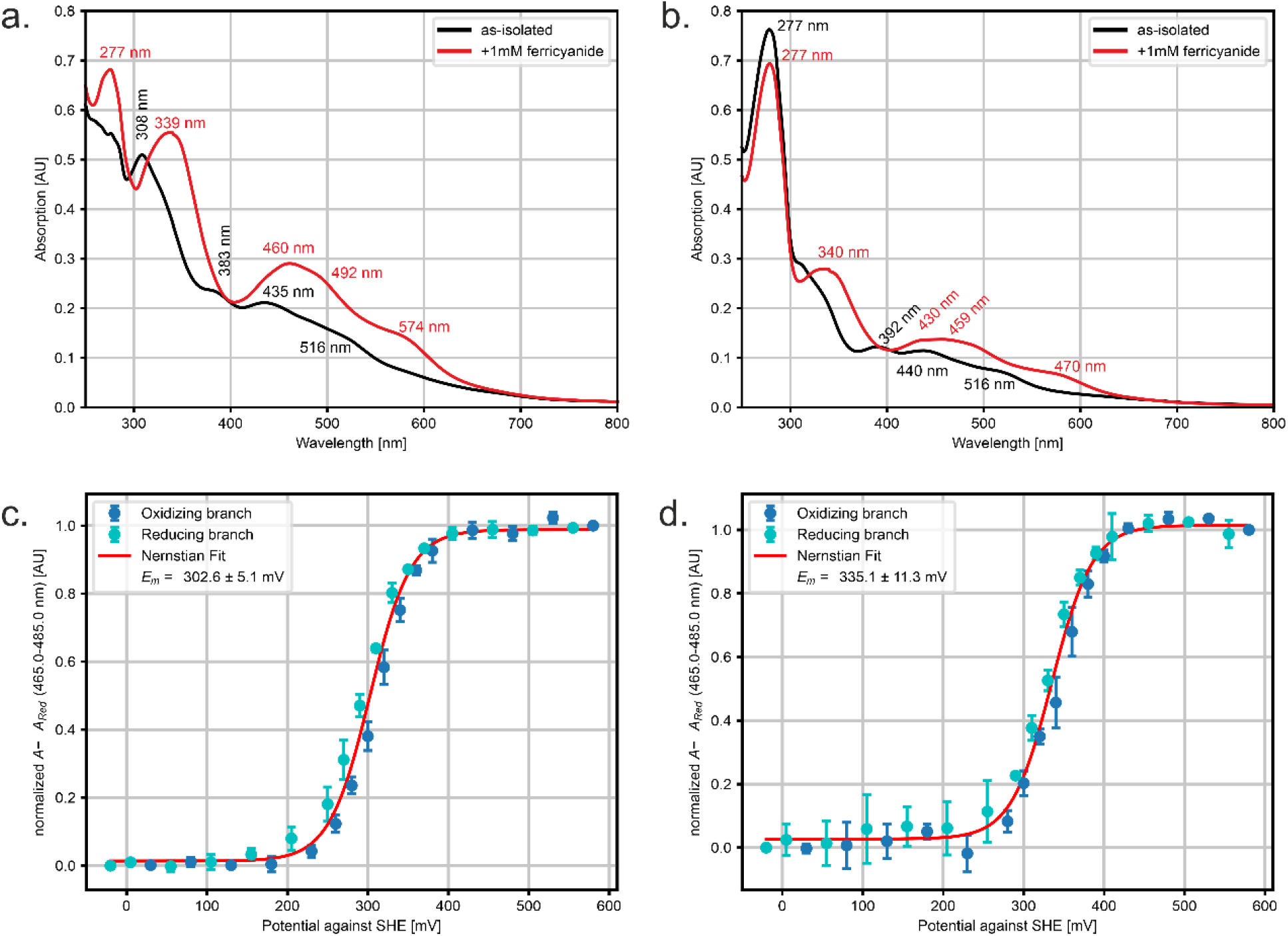
Top panels: UV/Vis spectra of Kuste4569 (**a**.) and Kustd1480 (**b**.) in the as-isolated state (black lines) and after oxidation with 1 mM potassium ferricyanide (red lines). Due to a lack of tryptophans in Kuste4569, the absorption peak at 277 nm is much lower than for Kustd1480. Peak wavelengths are indicated in nm. **Bottom panels:** spectroelectrochemical midpoint potential determination of Kuste4569 (**c**.) and Kustd1480 (**d**.). The oxidizing and reducing branches are shown in dark- and light blue, respectively, and the fit (which assumes a 1-electron transition) in red. All potentials are indicated *vs*. the Standard Hydrogen Electrode (SHE). Midpoint potentials of 303 ± 5 mV and 335 ± 11 mV were determined for Kuste4569 and Kustd1480, respectively.

Using an optically transparent thin-layer electrochemical (OTTLE) cell kept at fixed potentials with respect to an Ag/AgCl reference electrode, we measured spectra at potentials between -50 and +550 mV vs. Standard Hydrogen Electrode (SHE). For both proteins, these showed a single, clear transition in the averaged absorption between 465 and 485 nm which could be explained excellently with a single-electron redox transition (Figure 5c,d). Experiments were carried out in triplicates to obtain error bars on the individual absorption changes, which in turn were used to estimate the error in the observed midpoint potentials using bootstrap resampling. In this way, midpoint potentials of 303 ± 5 mV and 335 ± 14 mV could be determined for Kuste4569 and Kustd1480, respectively.

### Modeling of Rieske/cytochrome b complexes

Using AlphaFold3^11^, we prepared models of the complete Kuste4569-4573 (*i*.*e*., excluding Kuste4574) and Kustd1480-1485 complexes, assuming a dimeric state for the membrane-bound parts (the cytochromes *b*, SUIV-like and Rieske proteins) as well as for the associated *c-*type cytochromes on the positive membrane side, and a monomeric state for the proposed NAD(P) reductase domains on the negative membrane side (Figure 6, Figure 7). These initial models contained all heme *c* and *b* cofactors, as well as the iron-sulfur clusters in the Rieske proteins and FAD and NAD in the NAD(P) reductases. In both complexes, these latter domains showed a number of cysteine clusters tracing out a path between the Rieske/cytochrome *b* part and the active site of the NAD(P) reductases, likely binding [4Fe4S] and/or [3Fe4S] iron-sulfur clusters (Supplemental Figure S6), which were placed in these positions using homologous experimental structures as described in the methods section. Both models show high confidence in the core regions, with lower confidence values in loops on the outside of the protein.

**Figure 6.**
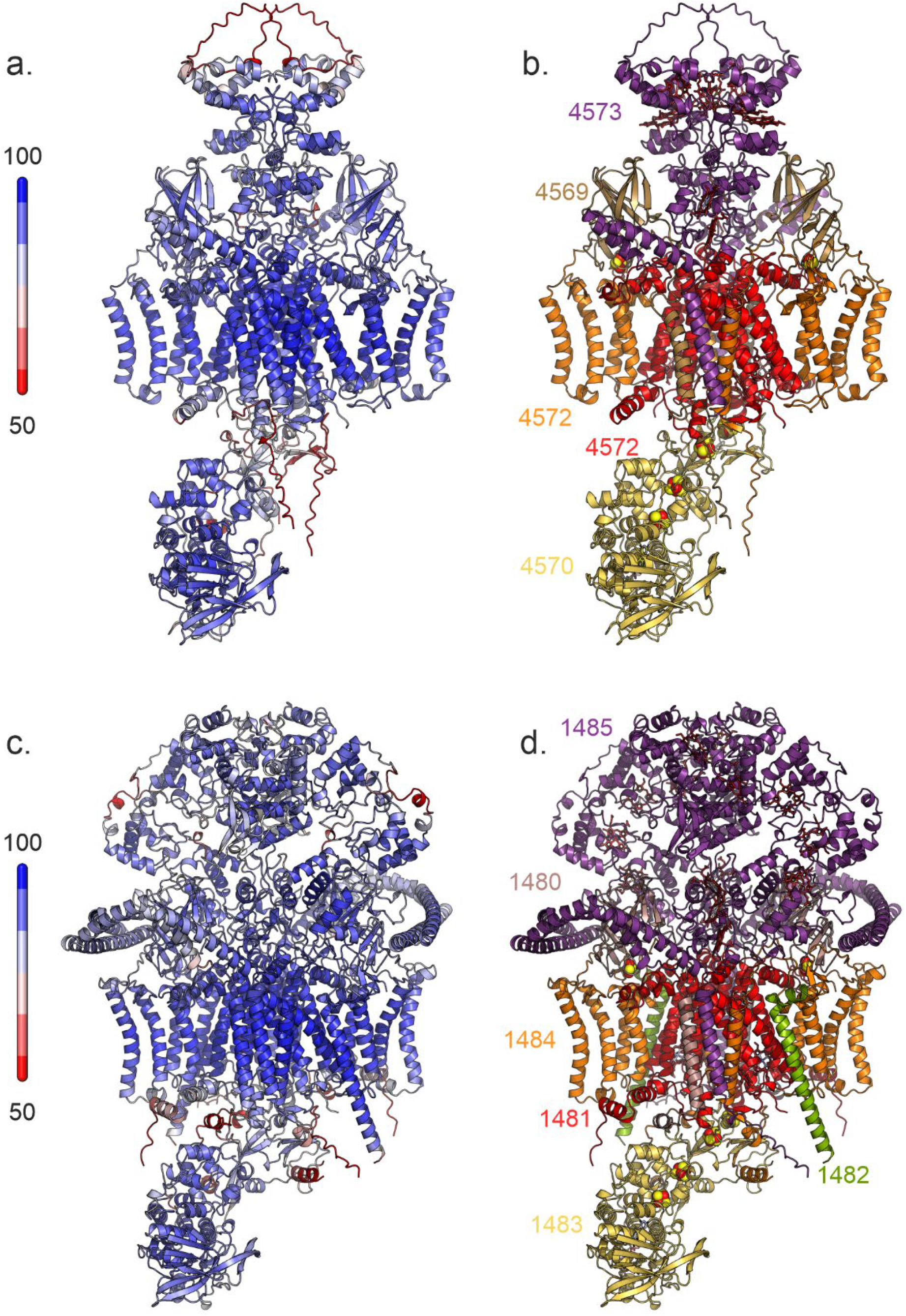
AlphaFold3^11^ models of the Kuste4569-4573 (a.,b.) and Kustd1480-1485 (c., d.) complexes. Panels a. and c. are colored according to plDDT, with red indicating a low confidence (plDDT ≤50) and blue indicating high confidence. Panels b. and d. are colored according to polypeptide chain.

**Figure 7.**
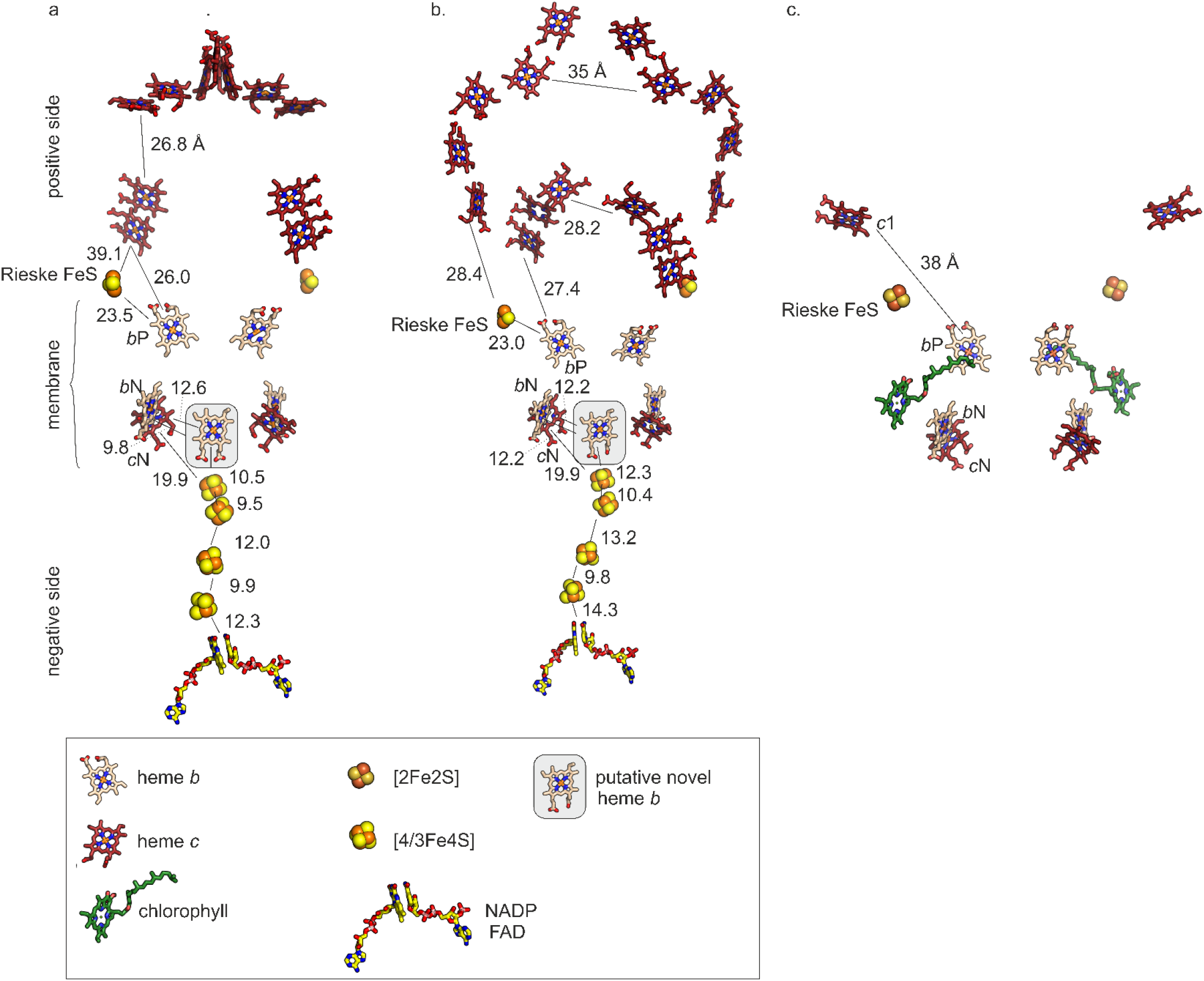
Cofactor positions and intercofactor distances in the models of **a**. Kuste4569-4573, **b**. Kustd1480-1485 and in the experimentally determined structure of the Rieske/*b*_*6*_*f* complex from spinach. The proposed novel *b*-type heme is indicated with a grey box. Edge-to-edge distances are indicated in Ångstrom.

The models suggest that the cytochrome *b* and subunit IV parts of both complexes are very similar to each other as well as to *b*_6_*f* complexes in general, except that the SUIV part has five instead of three transmembrane helices in both complexes. In crystal structures of *bc*_1_ complexes, the Rieske proteins can be observed in at least three conformations: ‘*b*’, ‘Int’ or ‘*c*_1_’, where the iron-sulfur cluster points either to the Q_P_ site in cytochrome *b*, to heme *c* of cytochrome *c*_1_ or takes on an intermediate position that is similar to the ‘*b*’ state ^38^. In our AlphaFold models, the Rieske proteins appear to be modeled in the ‘*b*’ or intermediate position in both complexes, with the [2Fe-2S] cluster pointing toward the quinone binding site. As expected from their sequences, the multiheme *c*-type cytochromes share only little similarity between the two complexes, though their transmembrane helices are in the same position in the complex and the two *c*-type hemes closest to the membrane are placed approximately in the same location. Kustd1485 is far larger than Kuste4573 and has long α-helices that wrap halfway around the complex including the Rieske protein Kustd1480.

The membrane hemes *b* and *c*_N_ (which are *c-*type hemes in the transmembrane part close to the negative side of the membrane) are positioned the same as expected for canonical Rieske/cytochrome *b* complexes (Figure 7). In contrast, the *c*-type hemes of the large multi-heme cytochromes show very different arrangements with minimum edge-to-edge cofactor distances to the membrane-bound cofactors of just 26-28 Å compared to 38 Å in the *Spinacia oleracea b*_6_*f* ^39^. The remaining *c*-type hemes are positioned in groups of two (Kuste4573, Kustd1485) and three (Kustd1485). Strikingly, a new *b*-type heme binding site at the interface of the cytochrome *b* dimer was predicted by AlphaFold3 for both complexes. In the Kuste4569-4573 complex model, this unexpected *b*-type heme is bound by the His43 residues of the two Kuste4571 monomers as the two iron-coordinating residues (Supplemental Figure S7), whereas its propionate groups are positioned in the vicinity of the Kuste4571 Arg247 side chains and the Kuste4570 Arg511 side chain. The iron-coordinating residues in the Kustd1480-1485 complex model are the two His62 residues from each Kustd1481 monomer, with the side chains of Arg266 and Lys267 from Kustd1481 and Arg550 from Kustd1483 positioned well to compensate the negative charge of the heme propionates.

## Discussion

Both Kuste4569 and Kustd1480 show high redox potentials of +303 and +335 mV, respectively. Relatively high (>200 mV) redox potentials would be expected given the presence of the hydrogen-bond donating serine residues in the cofactor’s second coordination sphere in both Rieske proteins (Ser127 in Kuste4569/Ser114 in Kustd1480) as well as Tyr129 in Kuste4569. Mutating these residues in the *S. cerevisiae* Rieske protein was reported to result in redox potential decreases of 130 mV for the serine mutant and 65 mV for the tyrosine mutant, with roughly an additive effect for the double mutant ^10^. These effects were of similar magnitude in mutants of the Rieske proteins from *B. taurus* ^40^ and *R. sphaeroides* ^*36*^. Accordingly, it is somewhat unexpected that Kustd1480 displays a potential even higher than that of Kuste4569, despite lacking one of these hydrogen-bonding residues. Possible reasons for this observation could include other differences in the second coordination sphere, surface electrostatics or pH dependence, but the discrepancy is not easily attributed to any single effect, and is likely a combination of several, more subtle influences. In any case, the measured potentials should be regarded with a degree of caution, since they were obtained for the soluble Rieske fragments in their isolated state, *i*.*e*. removed from the environment of their respective Rieske/cytochrome *b* complex. However, the measurements with or without complex tend correspond to within a few tens of millivolts, such as in the Rieske proteins from *R. sphaeroides*^41-44^ and from *S. oleracea*^45,46^, which perhaps is understandable given that Rieske proteins assume various positions and orientations in their parent complexes.

Nevertheless, the potentials of both Rieske proteins are higher than would be expected for a canonical, menaquinol-oxidizing Rieske/cytochrome *b* complex. In the Q cycle mechanism of conventional Rieske/cytochrome *b* complexes, the energy required for proton translocation is derived from the electron transfer from quinol to the Rieske protein, and corresponds to the potential difference between them. However, the number of protons that are translocated in this mechanism is fixed to exactly two per half-cycle, and the energy required depends mostly on the proton-motive force. Consistent with this, the redox potentials of Rieske proteins and quinone species tend to correlate well across different organisms and low-potential quinones are found together with Rieske proteins of lower potentials (<200 mV) ^5,7,43^. Indeed, the potentials of the Rieske protein and heme *b*_P_, *i*.*e*. the two redox centers to which the electrons are transferred first upon quinol oxidation and electron bifurcation, are typically symmetrical around the quinol potential (Figure 8)^43^. Thus, the observed high potentials of the Rieske proteins investigated here would suggest that the potential of heme *b*_P_ could be as low as -475 mV. Even accounting for the fact that the Rieske redox potentials might be somewhat different when the proteins are embedded in their parent complexes, this would allow electrons from heme *b*_P_ to be used for the reduction of NAD(P) as in the original proposal by Kartal *et al*.^1^, given its redox potential of ∼-320 mV, if this reaction is coupled to an energetically favorable reaction such as nitrite reduction by the electrons taking the high-potential pathway *via* the Rieske protein.

**Figure 8.**
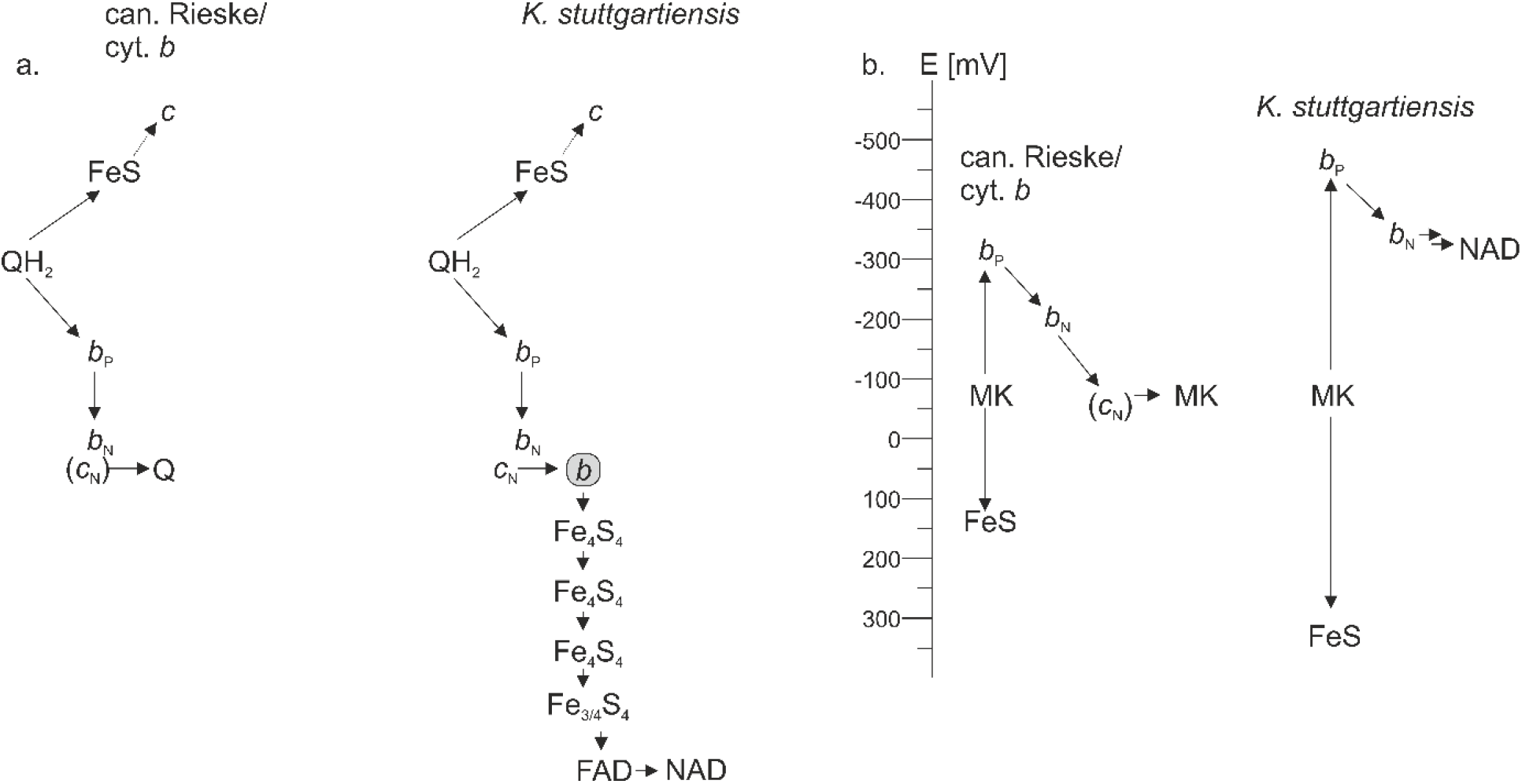
Electron transfer in Rieske/cyt. *b* complexes. **a.** electron transfer pathways in a canonical Rieske/cyt. *b* complex (left) and as proposed for the Kuste4569-4573 and Kustd1480-1485 complexes in *K. stuttgartiensis*. QH_2_: reduced lipoquinone, Q: oxidized lipoquinone, FeS: Rieske iron-sulfur protein. *b*_P_: positive-side heme *b, b*_N_: negative-side heme *b*, positive-side heme *b, c*_N_ negative-side heme *c, b*: putative novel heme *b*, Fe_4_S_4_ and Fe_3_/_4_S_4_: [4Fe4S] and [3Fe4S] iron-sulfur clusters, FAD: flavine adenine dinucleotide, NAD: NAD(P)+. **b**. Energetics of electron transfer. In canonical complexes (left), the potentials of the Rieske FeS cluster and the positive side heme *b* (*b*_P_) are symmetrical around the potential of the lipoquinone oxidized, in this case menaquinone (MK); the potentials shown here are taken from the analysis of the *H. modesticaldum* complex by Bergdoll and coworkers^43^. In the case of the unusual *K. stuttgartiensis* complexes (right), this symmetry would suggest a very low potential for the *b*_P_ heme, energetically allowing the reduction of NAD(P)+ to NAD(P)H.

However, to properly judge the viability of such a scheme one needs to consider the possible electron transfer pathways and redox potentials involved. To be transferred to the putative NAD(P) oxidoreductase subunits, electrons coming from heme *b*_P_ would first need to be transferred to heme *b*_N_. As shown in Figure 7, the edge-to-edge distances would allow efficient transfer between these two hemes, and if the trend of ∼130 mV redox potential difference between hemes *b*_P_ and *b*_N_ outlined by Bergdoll and coworkers ^43^ also apply to the two anammox complexes investigated here, potentials of -345 to -315 mV can be estimated for heme *b*_N_ (based on the measured Rieske potentials). Such low potentials are not unrealistic, given that there are Rieske/cytochrome *b* complexes in *Firmicutes* in which the *b*_P_ and *b*_N_ hemes display very low potentials of -360/-220 mV ^43^. From heme *b*_N_ onward, the electrons would only need a viable transfer pathway to the active site of the putative NAD(P) oxidoreductase.

The AlphaFold3 models of the complexes trace out just such a pathway (Figure 7), starting from the newly predicted, unexpected *b*-type heme at the center of the homodimer that could accept electrons from the hemes *b*_N_ or *c*_N_ from either monomer (Figure 7). The electrons could then be shuttled along the four [4/3Fe4S] clusters to the FAD cofactor with a maximum of 15 Å tunneling distance for each step. This would enable efficient electron transfer rates^47^, but even when not taking the novel *b*-type heme binding site into account, electron transfer from heme *c*_N_ to the first [4Fe4S] cluster could still be possible at 20 Å. The efficiency of NAD(P) reduction would depend on the redox potentials of hemes and iron-sulfur clusters involved, but, in theory, their potentials could cover the required range^48^,

The electron pathways on the positive side of the membrane (where the *c*-type cytochromes are located) are less obvious but, as seen with canonical *bc*_1_ and *b*_6_*f* complexes, the soluble domain of the Rieske protein will likely rotate to approach the closest *c*-type heme and reduce it. As mentioned in the results, the distance between hemes *b*_P_ and the *c*-type hemes are surprisingly low, when one considers that the energetically favorable short-circuit would be undesirable. Furthermore, the need for the large number of *c*-type hemes in the subunits on the positive side of the membrane is not entirely clear, but for Kuste4569-Kuste4573, some of these hemes could be involved in further electron transfer to the proposed nitrite-reducing Kuste4574. Unfortunately, no reasonable models of the complex including Kuste4574 were predicted by AlphaFold3, which is unfortunate since comigration of all subunits was observed experimentally^8^ indicating the formation of a stable complex. Electron transfer (*via* the novel heme *b* or not) to the NAD(P) oxidoreductase subunit would make it unlikely that the complexes perform quinone reduction at the negative membrane side as in the canonical Q-cycle. Indeed, sequence information did not unambiguously support the presence of functional Q_N_ sites as noted by the original proposers of the novel mechanism^1^. We will return to this issue later.

Estimates of the redox potentials of the *c*_N_ hemes (Figure 7) are more complicated to make. In the chloroplast *b*_*6*_*f* complex, the *c*_N_ heme’s redox potential is around +100 mV – the same as that of plastoquinone ^49 6^. This match in potential appears to follow a pattern, as it is also observed in organisms using menaquinone (here it must be noted that redox potential determination is complicated and the values are obtained with low confidence only) ^43,50^. Importantly, the largest influence on the redox potential of heme *c*_N_ is the second axial heme ligand. If this second ligand is a water, high potentials are observed for heme *c*_N_, while a glutamate as second ligand leads to low potentials (around -50 mV) are obtained ^43,51^. Despite a lack of experimental structures of *b*_*6*_*f* complexes and glutamate-coordinated hemes *c*_N_, clear EPR evidence for such a coordination exists ^51^. That glutamate coordination is indeed the major factor governing redox potential is supported by the observation that addition of the coordinating inhibitor NQNO results in a considerable decrease in heme *c*_N_ redox potential ^49^. Indeed, NQNO binding was observed directly in an experimental structure^52^). Analysis of a Phe-Tyr mutant in the equivalent position to glutamate second ligand in low potential hemes *c*_N_ further supports this residues role in redox potential tuning^53^.

In our models of Kustd1480-1485 and Kuste4569-4574, the hemes *c*_N_ are indeed ligated by glutamate residues, which suggests a redox potential of at least around -50 mV for these hemes. Such a redox potential would still be ∼250 mV higher than the likely potentials of the rest of the cofactors in the electron transport chain. A lower potential would, however, appear reasonable if the electrons were not shuttled towards menaquinone-but to NAD(P)+ reduction. For the Phe-Tyr mutant^53^ mentioned above, a potential of -200 mV was observed, showing at least even lower potentials to be possible ^53^. However, as the edge-to-edge cofactor distances are short if the novel *b*-type heme is present, electron transfer should still be possible in the case of a ‘regular’ heme *c*_N_ potential, even if that would require a step that is energetically uphill by ∼ 250 mV.

A far larger problem these complexes would have to solve would be the prevention of menaquinone reduction at the membrane’s negative side. Possibly, having a compromised quinone binding site as opposed to canonical *b*_*6*_*f* complexes would already suffice. However, this last electron transfer step in the canonical Q cycle mechanism is expected to be energetically highly favorable. In our complex models, the *c*_N_ hemes appear only somewhat less accessible than their counterparts in structures of canonical *b*_*6*_*f* complexes and could allow close approach by a quinone. Moreover, a histidine is found very close to both the heme and glutamate in the two complexes. This histidine could assist a proton-coupled electron transfer as in quinone reduction but would have no clear function in a simple heme-to-heme electron transfer step. One could argue that, given the presence of the novel heme *b*, NAD(P)H could be used to refill the quinol pool through reduction close to heme *c*_N_. This, however, would mean incurring an enormous energy loss, and since *K. stuttgartiensis* expresses NADH dehydrogenases that can couple this reaction with proton (or sodium) translocation more efficiently^8^ appears unlikely. Thus, our experimental redox potentials and the models appear to be most consistent with the original proposal ^1^ that these complexes serve to produce NAD(P)H, and that they do so by coupling NAD(P) reduction to another, more favorable process such as nitrite reduction.

Returning to the Rieske proteins themselves, the observed flexibility in the Kustd1480 cofactor binding domain and C-terminal loop appears puzzling. Typically, electron transfer proteins display little conformational flexibility to minimize reorganization energy upon redox state changes. Possibly, the observed flexibility is an artefact caused by removing the Rieske protein from its natural surroundings in the complex. However, it is also possible that it is part of an induced-fit-type mechanism that allows the Rieske protein to adapt to the various positions and orientations inside the complex it needs to assume.

In summary, the Rieske iron-sulfur cluster proteins investigated here display high redox potentials, which can at least in part be explained by their crystal structures. These high redox potentials align very well with the proposed, novel function - NAD(P)H generation-of the parent complexes of these proteins. Finally, models of these complexes provide a basis for the explanation and further investigation of the mechanism of NAD(P)H generation (and possibly nitrite reduction) by the unusual Rieske/cytochrome *b* complexes in anammox organisms.

## Acknowledgements

The authors thank the staff of the Swiss Light Source (Villigen CH) for their support and facilities, Ilme Schlichting and Miroslaw Tarnawski for expert help with crystallization, crystal harvesting and data collection, and Melanie Müller for peptide map fingerprinting. Furthermore, TRMB is very grateful to Ilme Schlichting and KP to Arne Möller for continuous support over many years. This work was funded by the Max Planck Society as well as by DFG Grant 514060533 to KP and TRMB.

## Data availability statement

Crystal structures and diffraction data were deposited in the Protein Data Bank ^54^ as entries 9rk3 and 9rk4. AlphaFold3 models will be made available publicly *via* the Zenodo repository upon eventual acceptance of the manuscript.

