## Supplemental Information for "Rieske Iron-Sulfur Cluster Proteins from an Anaerobic Ammonium Oxidizer Suggest Unusual Energetics in their Parent Rieske/cytochrome *b* complexes"

*Correction of diffraction intensities for a lattice translocation defect in the crystals of Kuste4569*

After phasing of the Kuste4569 data, refinement failed to bring down the R-factors. Moreover, additional density not explained by the MR solution was seen in the electron density maps, which looked like additional proteins molecules. However, building protein molecules into this “ghost density” would have caused severe clashes.

A possible explanation for such observations is a lattice translocation defect. Indeed, inspection of the intensities revealed a strong modulation along the *h* direction in reciprocal space with a periodicity of 5 lattice planes (Supplemental Figure S1), changing sign with every lattice plane along *k*, suggesting a noncrystallographic translation of around 0.2 along x and around 0.5 along y. Indeed, the strongest off-origin peak in the Patterson map was at 0.223,0.5,0. However, this translation was already accounted for by the molecular replacement solution.

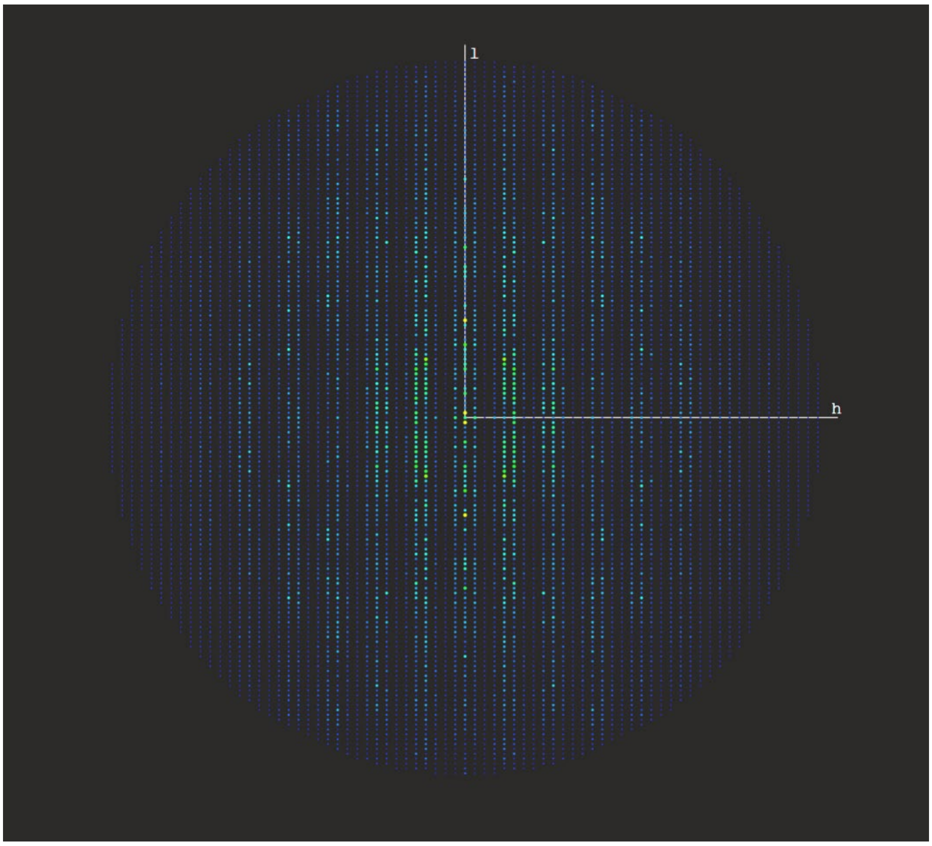


**Supplemental Figure S1.** *k*=7 plane of reciprocal space; a strong modulation of the intensities is seen in the *h* direction. This is, however, due to noncrystallographic translation symmetry and does not explain the problems with structure refinement.

Importantly, the next highest Patterson peak, which was not accounted for in the MR solution was found at (0.439,0,0). Translating molecules by this vector perfectly placed them in the ghost density.

These are clear indications of a lattice translocation defect [1]. Lattice translocation defects can occur when the crystal layers have more than one possible way of stacking, allowing for part of the layers to be shifted in one plane relative to the others (Supplemental Figure 2). The displaced molecules are then visible in the electron density in the form of a 'ghost molecule' that was shifted relative to the main structure. Moreover, it leads to a strong off-origin Patterson peak.

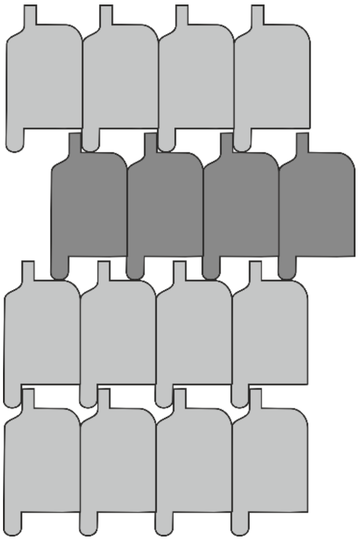

**Supplemental Figure S2.** Schematic example of a lattice translocation defect, redrawn from [1]. The dark grey layer of molecules is shifted with respect to the other, light grey, layers of molecules.

A method of correcting for these kinds of lattice defects was described by [2]: Suppose that there are *N* layers in a crystal which are offset from the other 'normal' layers *M* by the displacement vector $\boldsymbol{t_d}$, then the X-rays phase difference of the X-rays diffracted by those layers with respect to the normal ones will be $e^{2\pi i \boldsymbol{h t_d}}$, where $\boldsymbol{h}$ is the reciprocal vector (*h*, *k*, *l*). Using this relationship, the total structure factor $\boldsymbol{F}_{total}$ can be described as follows, where $\boldsymbol{F}_{unit}$is the structure factor for a unit cell:

$$\boldsymbol{F}_{total} = [M + Ne^{2\pi i \boldsymbol{h t_d}}]\, \boldsymbol{F}_{unit}$$

The number of layers can also be expressed in relative terms as their fraction of total crystal layers, $\kappa$ and $1\text{-}\kappa$. Redefining $\boldsymbol{F}_{total}$ as the observed structure factor of a unit cell with translation defects and F_unit as the structure factor of a unit cell without translation defects, one obtains

$$\boldsymbol{F}_{total} = [\kappa + (1-\kappa)e^{2\pi i \boldsymbol{h t_d}}]\, \boldsymbol{F}_{unit}$$

Or, in terms of the actual observed intensity measurements

$$I_{total} = \left|\kappa + (1-\kappa)e^{2\pi i \boldsymbol{h t_d}}\right|^2 I_{unit} = f I_{unit}$$

with

$$f = A + B\, cos(2\pi \boldsymbol{h t_d})$$

$$A = \kappa^2 - 2\kappa + 1$$

$$B = 2\kappa(1-\kappa)$$

Thus, the total observed intensity can be corrected for the effect of the lattice translocation by dividing it by $f$, which is a function of the fraction of displaced molecules $\kappa$ and the displacement vector $\boldsymbol{t_d}$. The latter can be obtained from the Patterson map, and the former can be determined using a trial-and-error method, by trying to find a value for $\kappa$ that minimizes the height of the Patterson peak corresponding to $\boldsymbol{t_d}$.

To that end, we wrote a Python script that reads and writes MTZ files using the SFTOOLS program of the CCP4 suite and that performs the required corrections for a preset value of $\kappa$. After a quick, coarse screen, we prepared corrected datasets using $\boldsymbol{t_d} = (0.439, 0.0, 0.0)$ as determined from the Patterson map and values for $\kappa$ of 0.10, 0.11, 0.12, 0.13, 0.14 and 0.15. The best available structure was then refined against these datasets, and the best improvement for both $R_{work}$ and $R_{free}$ was observed for $\kappa$ =0.13. Indeed, at this value for $\kappa$, the Patterson peak caused by the translocation defect is almost completely absent from the Patterson map (Supplemental Figure S3), and the electron density map showed almost no evidence of "ghost molecules".

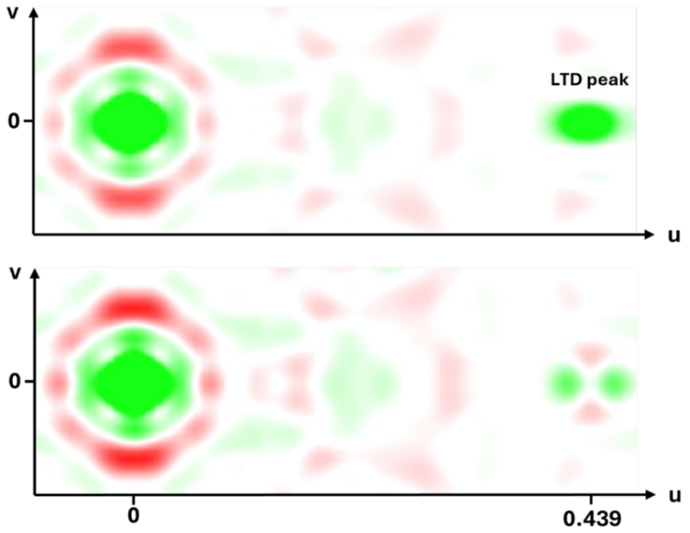


**Supplemental Figure S3.** Correction for lattice translocation defect. Top: Patterson section *w=0* before, and bottom: after correction of the intensities. The peak indicating a lattice translocation defect ("LTD peak") is almost completely gone. The color scale runs from -9 σ (red) to +9 σ (green).

As can be seen in Supplemental Figure S3, the Patterson peak caused by the defect is not spherical in nature but elongated along the *u*-axis. One reason for this could be that the shifted layers are not translated by one exact distance but have some "play" in their displacement. The intensity correction function does not account for this, which is why the corrected Patterson map still shows some positive and negative peaks around the position of the original LTD peak.

Nevertheless, after the correction, rebuilding and refinement could be continued as with normal data unaffected by dislocation defects.

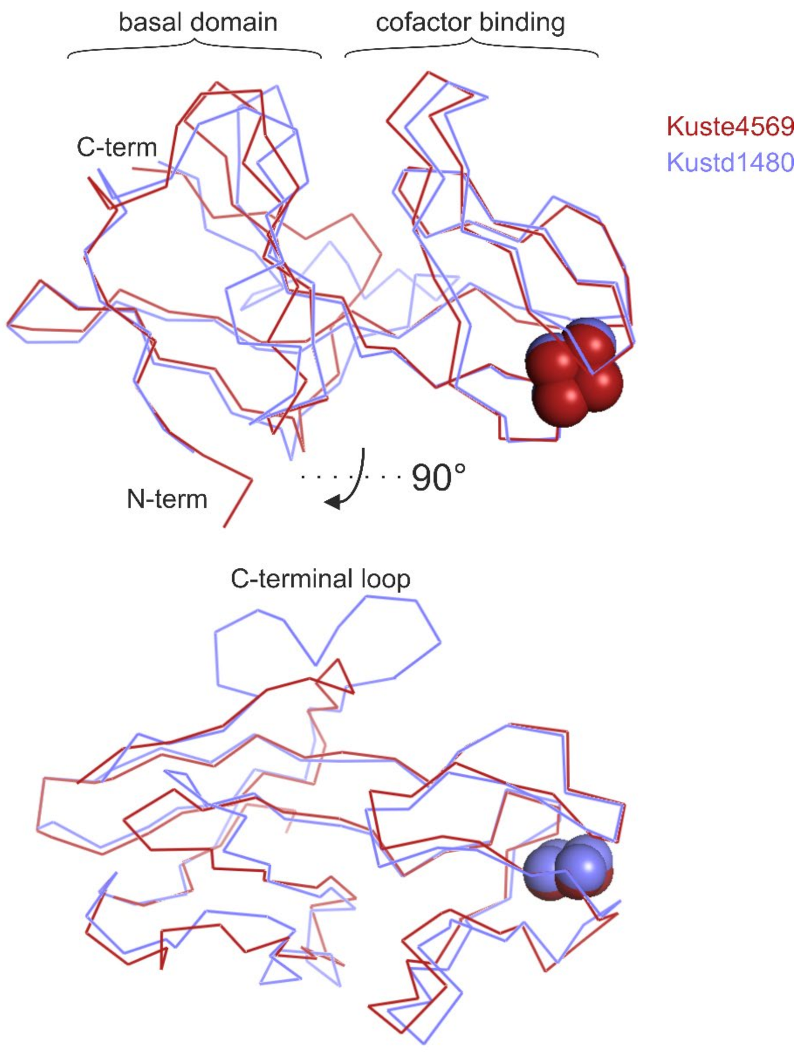


**Supplemental Figure S4.** Superposition of the crystal structures of Kuste4569 (red) and Kustd1480 (blue). The polypeptide chains are shown as a Cα trace, the cofactors as spheres.

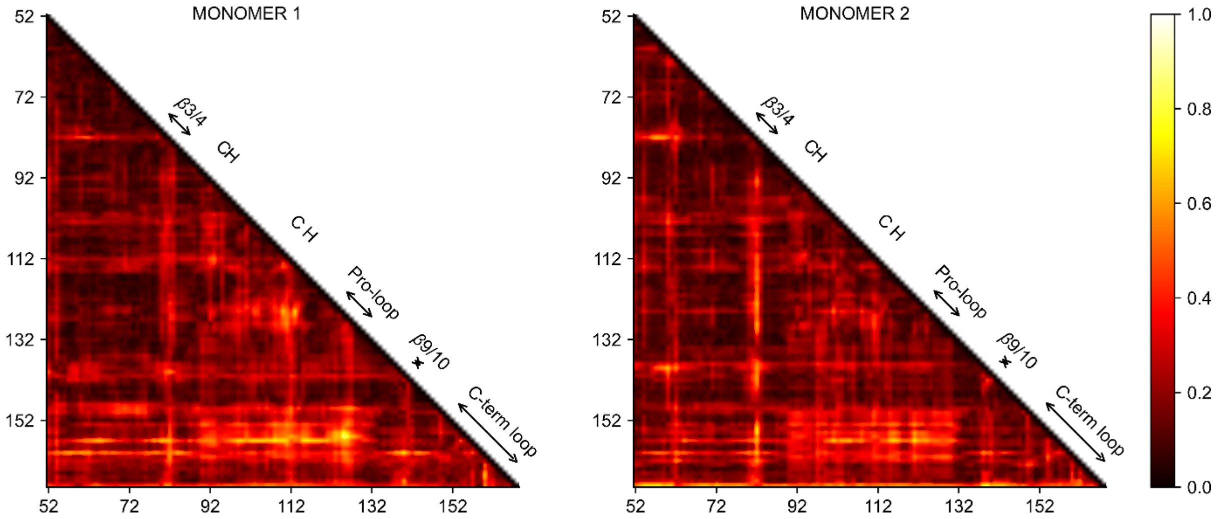


**Supplemental Figure S5.** RMSD of Cα- Cα distances from ensemble refinement for the two monomers in the Kustd1480 asymmetric unit. The color scale runs from 0 (black) to 1 Å (white). The highest flexibility is found in the C-terminal loop with respect to the cofactor-binding domain.

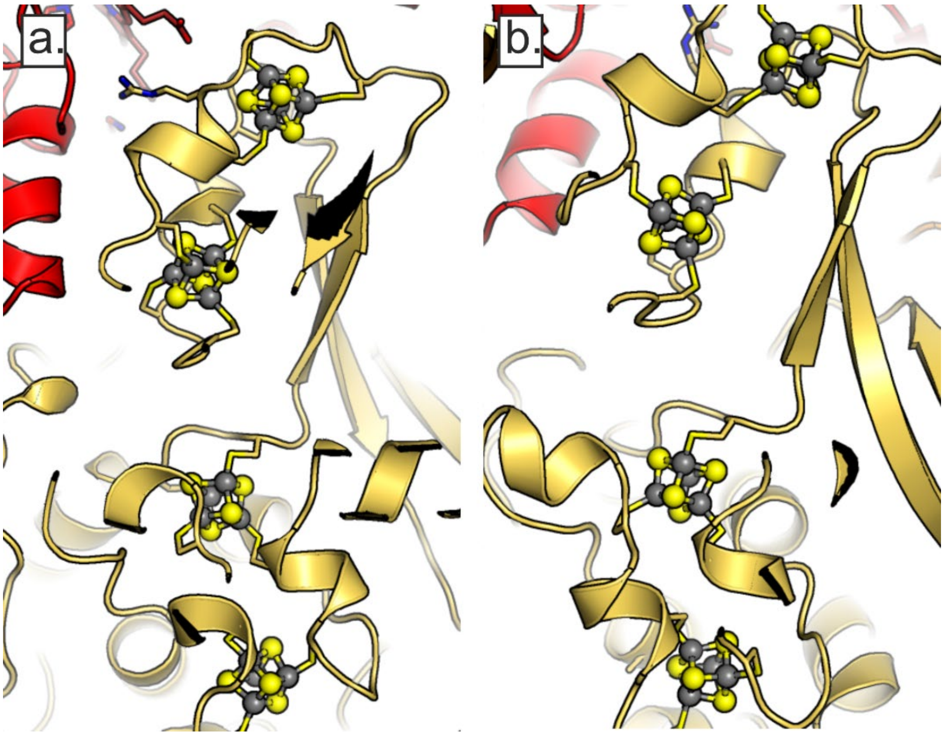


**Supplemental Figure S6.** Iron-sulfur clusters (ball-and-stick representation) bound by cysteine residues (stick representation) in the putative NAD(P)-oxidoreductase domains of the AlphaFold3 models of a. Kuste469-4573 and b. Kustd1480-1484

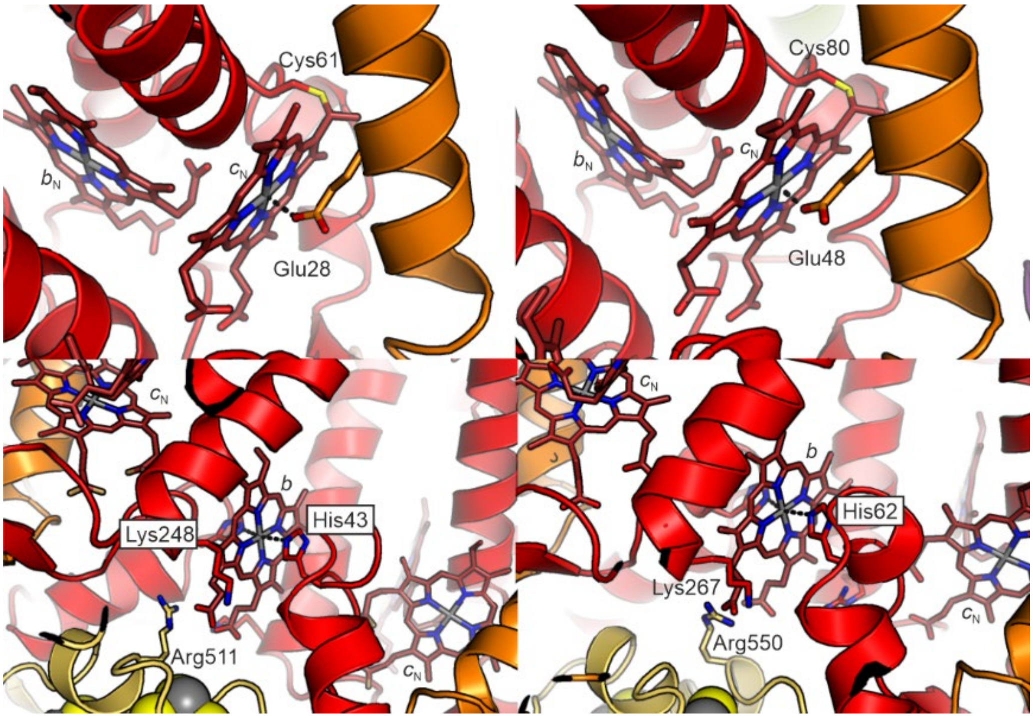


**Supplemental Figure S7: Special hemes in the Rieske/cyt *b* complex models.** (A) Hemes $c_n$ in the complexes of Kuste4569-4573 (left) and Kustd1480-1485 (right). The glutamates that were modeled as axial ligands are indicated. (B) Novel hemes *b* at the interface between the cytochrome *b* monomers. Axial coordination is symmetrical between the two monomers. Both hemes are positioned directly at the interface to the NAD(P) oxidoreductases Kustd1483 and Kuste4570 (yellow).
